# A simple image processing pipeline to sharpen topology maps in multi-wavelength interference microscopy

**DOI:** 10.1101/2023.01.12.523706

**Authors:** Peter W. Tinning, Jana K. Schniete, Ross Scrimgeour, Lisa S. Kölln, Liam M. Rooney, Trevor J. Bushell, Gail McConnell

**Author notes:** Joint first authors (Peter W. Tinning, Jana K. Schniete, Ross Scrimgeour).

## Abstract

Multi-wavelength standing wave (SW) microscopy and interference reflection microscopy (IRM) are powerful techniques that use optical interference to study topographical structure. However, the use of more than two wavelengths to image the complex cell surface results in complicated topographical maps and it can be difficult to resolve the three-dimensional contours. We present a simple image processing method to reduce the thickness and spacing of antinodal fringes in multi-wavelength interference microscopy by up to a factor of two to produce clearer and more precise topographical maps of cellular structures. We first demonstrate this improvement using model non-biological specimens, and we subsequently demonstrate the benefit of our method for reducing the ambiguity of surface topography and revealing obscured features in live and fixed cell specimens.

---

Interference-based microscopy techniques are a proven tool in the study of internal and external cellular structures. One of the most prominent interference-based microscopy techniques is standing wave (SW) microscopy, which was first demonstrated by Lanni et al. (1986) [1]. The image contrast in SW microscopy arises from an interference effect by one of two main methods; either the specimen is illuminated from opposite directions with two beams [1] or the specimen is placed in contact with a mirror [1,2]. The optical interference pattern that results from the SW is used to excite fluorescence from the specimen. In SW microscopy the antinodal fringe thickness is 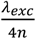 [1,3,4], where λ_exc_ is the excitation wavelength used and n is the refractive index of the specimen. Antinodal planes are axially separated by 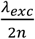 in air, this results in a sampling density that is approximately 50%. SW microscopy is compatible with imaging of fixed and live cells using confocal and widefield microscopy [1–4].

The recently developed TartanSW method is a multi-wavelength version of SW microscopy which uses multiple wavelengths that are close to the peak excitation wavelength of the fluorophore to excite the specimen. This increases the axial sampling from approximately 50% observed for single-wavelength SW imaging to up to 98%, but there exists considerable overlap in the axial positions of the antinodes [5]. This results in complex and low contrast images, and the cell topography can be difficult to extract from the multi-colour datasets.

Another technique that makes use of the principle of optical interference to obtain axial super-resolution is the label-free method interference reflection microscopy (IRM). Since the development of the technique in 1964, this method has been applied to a variety of live and fixed cell specimens for the observation of cellular features with an axial resolution which exceeds that possible with widefield and confocal microscopy [6,7]. As in SW microscopy, the contrast in IRM arises from the interference of light waves. However, IRM relies on reflected waves from different refractive index boundaries at the specimen plane, and this method is used to produce topographical images of unstained cellular specimens [7]. The antinodal plane thickness in IRM is given by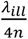 where λ_ill_ is the wavelength of illumination, and n is the refractive index of the specimen [7,8]. Antinodal interference fringes are axially separated by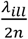. The numerical aperture (NA) of the imaging objective in IRM dictates the depth of field, which determines the number of interference fringes that can be detected [7,9].

It has been recently shown that IRM can make use of multiple illumination wavelengths to gain insights into the motility of bacterial cells [10]. The use of additional wavelengths enabled more precise visualisation of the position of the cell membrane relative to the glass substrate. However, there existed considerable overlap in the fringe pattern which complicated the interpretation of gliding motility.

We present a simple image processing method to reduce the thickness and spacing of antinodal fringes in TartanSW microscopy and multi-wavelength IRM by up to a factor of two, to produce a sharper and more precise topographical map of cellular structures. We use a difference operation to identify the spatial overlap in antinodal fringes in SW and IRM images in different imaging channels and hence improve the axial sampling precision of the antinodal fringes. A difference operation is used instead of a simple subtraction to avoid the generation of negative intensity values.

We first performed three-wavelength TartanSW imaging of a model specimen and we used these data to test the method. A 30 mm focal length plano-convex lens (Edmund Optics) was coated with a solution of 0.01% (w/v) poly-L-lysine in H_2_O (Sigma Aldrich) for 45 – 60 minutes, and then washed in deionized H_2_O and blow dried. The curved side of the specimen was submerged overnight in a 30 μM solution of DiI (Invitrogen) and dimethyl sulfoxide (Sigma). The specimen was then washed again in deionized H_2_O prior to imaging.

The lens specimen was placed with the curved surface in contact with a plane aluminium reflector (TFA-20C03-10, Laser 2000) and TartanSW imaging was performed using an upright widefield epifluorescence microscope (BX50, Olympus) with a 10x/0.4 dry objective lens (UPlanSApo, Olympus). Illumination was provided sequentially from 490 nm, 525 nm and 550 nm light emitting diodes (LEDs) (pE-4000, CoolLED). Emitted fluorescence was collected using a CMOS camera (ORCA-Flash 4.0LT, Hamamatsu) which was used with a 2.99x magnification camera port between microscope and camera. The LEDs and camera were synchronized and triggered using WinFluor software [11] with a 100 ms exposure time for each LED.

Using Fiji [12], a multi-colour merge of the individual SW images from each excitation channel was performed to create the TartanSW image. This was followed by a difference operation using the image calculator function in Fiji, firstly with the difference between images obtained with 525 nm and 490 nm illumination, and then with the difference between images obtained with 550 nm and 525 nm illumination. A two-colour merge of the individual difference images was then performed in Fiji. To quantify the full width at half-maximum (FWHM) antinodal fringe thickness and antinodal spacing in both TartanSW and the difference images a radially averaged line intensity plot was obtained for each channel of both datasets using the MATLAB script published previously [13].

The data from the imaged lens specimen are shown in Fig. 1. Fig. 1A shows the TartanSW image with a false-colour merge, using images obtained with excitation wavelengths of 490 nm, 525 nm, and 550 nm as blue, green, and red respectively. The radially averaged line intensity plot of the TartanSW image is shown in Fig. 1B. Fig. 1C shows the result of the image difference operation, with the difference between 525 nm and 490 nm shown in green and the difference between 550 nm and 525 nm displayed in magenta. The radially averaged line intensity plot from this difference operation is shown in 1D.

**Fig. 1.**
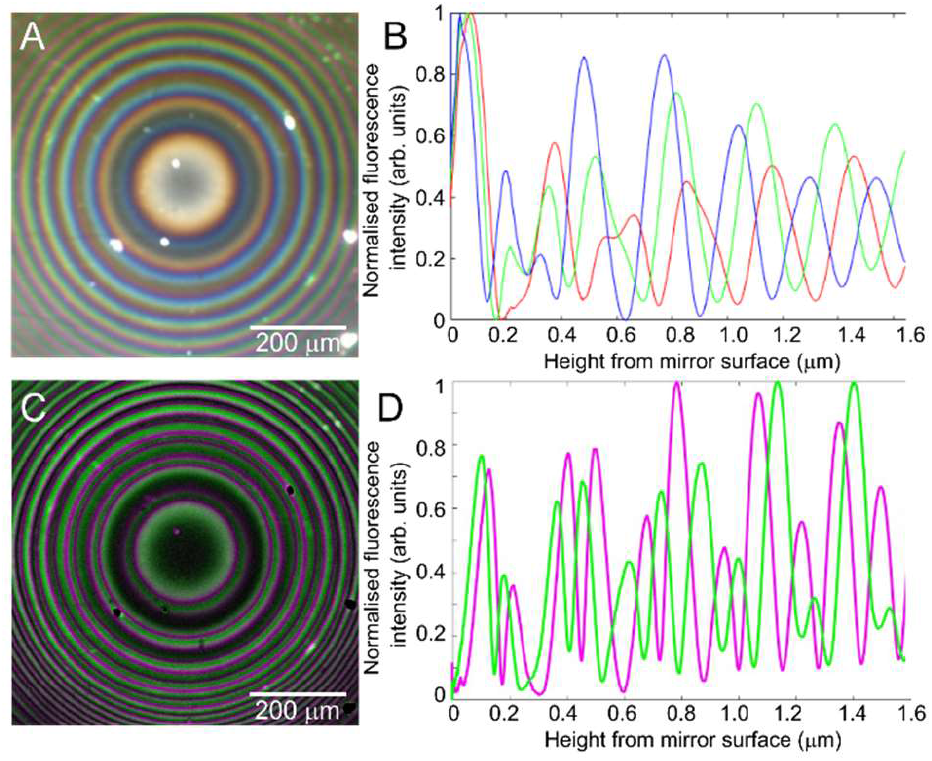
A) TartanSW image of a lens specimen prepared with a monolayer of DiI on the curved surface. Sequential excitation wavelengths of 490 nm, 525 nm and 550 nm were applied, and the resultant images are false colored in blue, green, and red look-up tables respectively. B) Radially averaged line intensity plot showing fluorescence intensities of individual channels of the TartanSW image in (1A) for 490 nm (blue), 525 nm (green) and 550 nm (red) excitation wavelengths with respect to the distance from the mirror surface. C). A difference image of |525 nm – 490 nm| (green) and |550 nm – 525 nm| (magenta) of the same individual channels used to create (1A). D) Radially averaged line intensity plot showing fluorescence intensities of individual channels of the difference image in (1A) for |525 nm – 490 nm| (green) and |550 nm -525 nm| (magenta).

Table 1 shows the experimental values for FWHM antinodal fringe thickness and antinodal fringe spacing for each of the illumination wavelengths used for TartanSW imaging of the lens specimen shown in Fig. 1B, together with the theoretical values for these parameters.

**Table 1.** Measured and theoretical values of antinodal fringe thickness at FWHM and antinodal fringe spacing for TartanSW imaging of a *f*=30 mm lens specimen.

| $\lambda$ (nm) | FWHM antinodal fringe thickness (nm) | | Antinodal fringe spacing (nm) | |
| --- | --- | --- | --- | --- |
|  | Theory | Measured | Theory | Measured |
| 490 | 122 | 123.3 ± 3.2 | 244 | 245.3 ± 8.6 |
| 525 | 131 | 126.9 ± 1.8 | 262 | 267.4 ± 10.7 |
| 550 | 137 | 134.0 ± 4.4 | 264 | 275.0 ± 3.8 |

Table 2 shows the measured results for the FWHM antinodal fringe thicknesses and average antinodal fringe spacings for difference imaging obtained with illumination wavelengths of |525 nm – 490 nm| and |550 nm -525 nm| over a height of 1.6 µm from the mirror surface, extracted from Fig. 1D. Theoretical values shown in Tables 1 and 2 are calculated using 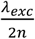 or from a simulation of the difference operation presented in Supplement 1. We measure up to a two-fold reduction in both FWHM antinodal fringe thickness and antinodal fringe spacing in the difference image compared to the TartanSW data. This reduction in fringe thickness facilitates an improvement in antinodal fringe sampling precision of the same factor. We also note an improvement in image contrast in the higher order antinodal fringes.

**Table 2.** Measured and average theoretical antinodal fringe thickness at FWHM and antinodal fringe spacing after applying the difference operation to TartanSW images of a *f*=30 mm lens specimen.

| Difference in $\lambda$ (nm) | FWHM antinodal fringe thickness (nm) | | Antinodal fringe spacing (nm) | |
| --- | --- | --- | --- | --- |
|  | Theory | Measured | Theory | Measured |
| 525 nm – 490 nm | 71.8 | 64.9 ± 5.8 | 123.2 | 129.0 ± 9.8 |
| 550 nm – 525 nm | 75.6 | 71.6 ± 3.2 | 131.7 | 136.7 ± 11.2 |

Next, we applied the difference operation to fluorescence images of live and fixed cell specimens obtained using widefield and confocal point scanning illumination. Live MCF-7 cells were labelled with the lipophilic membrane dye DiI and plated onto first surface reflectors (Laser2000) using the previously reported protocol [5]. The mirrors were submerged in 4 % BSA + PBS and imaged using the same protocol and equipment as described above with the exception that a 40x/0.8 water dipping objective lens (LUMPLFLN, Olympus) was used for cell imaging.

3T3 cells were plated onto the same type of mirror and fixed using 4 % paraformaldehyde (PFA) before labelling with rhodamine phalloidin (Invitrogen) using the method described previously [5]. Specimens were mounted in PBS and imaged using a confocal microscope (SP5, Leica) with a 40x/0.8 water dipping objective (HCX APO LUV-I, Leica). Excitation was provided sequentially using 488 nm, 514 nm and 543 nm laser lines with a scan speed of 100 Hz, with line averaging of 8 and an image size of 2048 pixels and no digital zoom. For each wavelength, fluorescence was detected between 550 – 650 nm. All raw image data were opened in Fiji. TartanSW images were created as reported previously [5] and the difference operation was applied using the same method as for the lens specimen.

TartanSW images of both live and fixed cell specimen are presented in Fig. 2. As shown in Figs. 2A & 2C, signals from higher order fringes appear with low contrast in the images, making it difficult to interpret the cell structure. Some adjustment of the gamma scale can be used to slightly improve this, but it is difficult to improve the colour specificity while avoiding saturation of the image. The application of the difference operation, evidenced by Figs. 2B and 2D with digitally zoomed regions of interest, improves the sharpness of the cell structure. This is evident in the TartanSW images of rhodamine phalloidin labelled F-actin. Our method selectively removed signals originating from low order fringes which overlap for the different wavelengths. The difference method removed saturated signals from the raw TartanSW images that originate from dense cell regions close to the mirror surface (see Fig 2C). As a result, actin structures at the basal cell membrane are more clearly visible (see Fig. 2D).

**Fig. 2.**
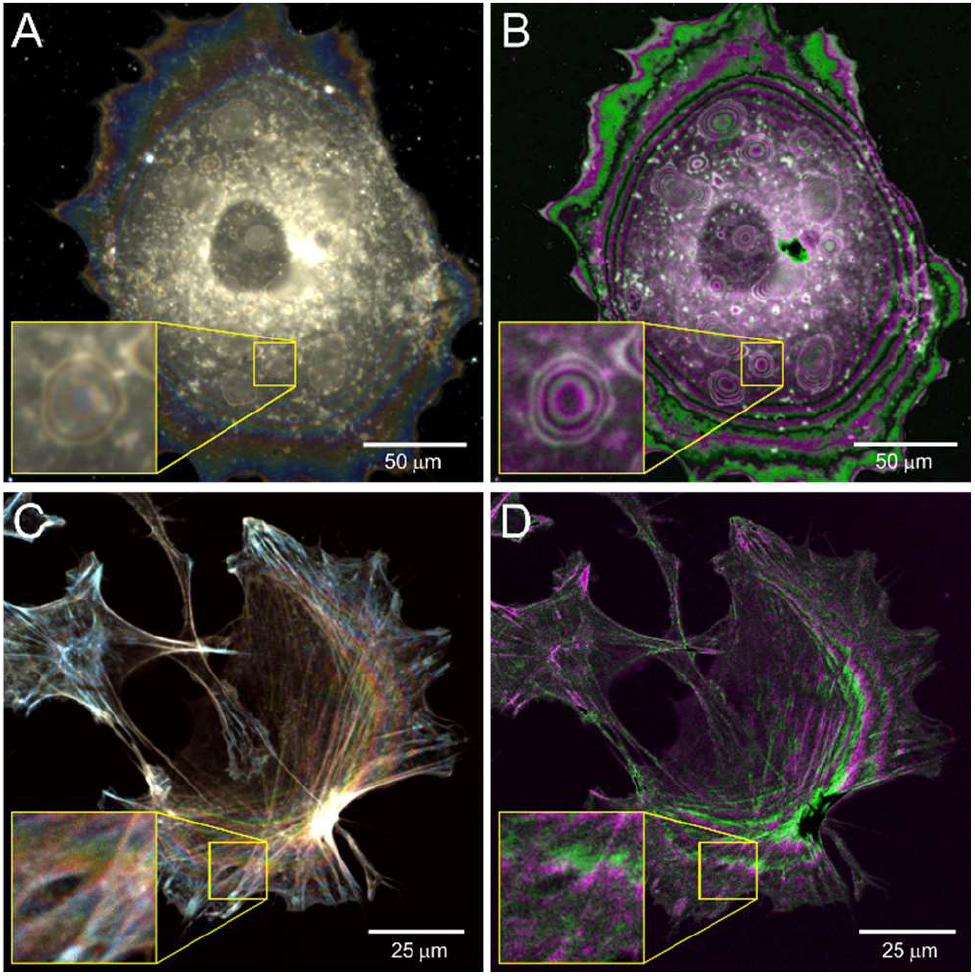
A) Widefield TartanSW image of live MCF-7 cells stained with DiI. B) Difference image of (2A) with |525 nm – 490 nm| in green and |550 nm – 525 nm| channel in magenta. C) Confocal TartanSW image of fixed 3T3 cells labelled with Rhodamine Phalloidin. D) Difference image of (2C) with |525 nm – 490 nm| in green and |550 nm – 525 nm| channel in magenta. Yellow boxes show regions of interest with a digitally expanded zoom of these areas for each image.

Time lapse widefield imaging using the difference operation to observe changes in cell topography was also performed. Live MCF-7 cells labelled with the membrane stain DiI were imaged using the same widefield microscope setup used to image the lens specimen, with 100 ms exposure time at 20 s intervals over a total period of 16 minutes, and the difference operation applied as reported. Visualization 1 shows an example dataset, where small changes in the cell topography are revealed as a shift in the position of the high contrast antinodal fringes.

We also applied the difference operation to images obtained with multi-wavelength IRM. Again, we first performed imaging of a 72 mm focal length plano-convex lens placed upon a microscope coverslip (631-0153, VWR). Since the contrast in IRM arises from reflection, no fluorescent stain was required. Multi-wavelength IRM was carried out using the same equipment and method as described by Rooney *et al*. [10]. The lens specimen was imaged using a confocal microscope set in reflection mode with a 10x/0.3 UPLANFL lens (Olympus, Japan) using 488 nm and 514 nm lasers, and the difference method was applied. Data are shown in Fig. 3A.

Fixed MeT-5A cells (ATCC, CRL-9444) were also imaged using multi-wavelength IRM to evaluate the value of the difference operation for the study of more complex structures. Cells were plated on poly-L-lysine-coated coverslips 24 hours prior to fixation in 4% PFA. The cells were mounted in ProLong Diamond Antifade mountant (Invitrogen) (*n*=1.46). The cell specimen was imaged with a confocal microscope (SP5, Leica), using a 20x/0.7 objective (506513, Leica), and a 488 nm, 514 nm or 633 nm laser line for illumination. A 10-frame average was taken, and the difference operation was performed. Data are shown in Fig. 3B.

**Fig. 3.**
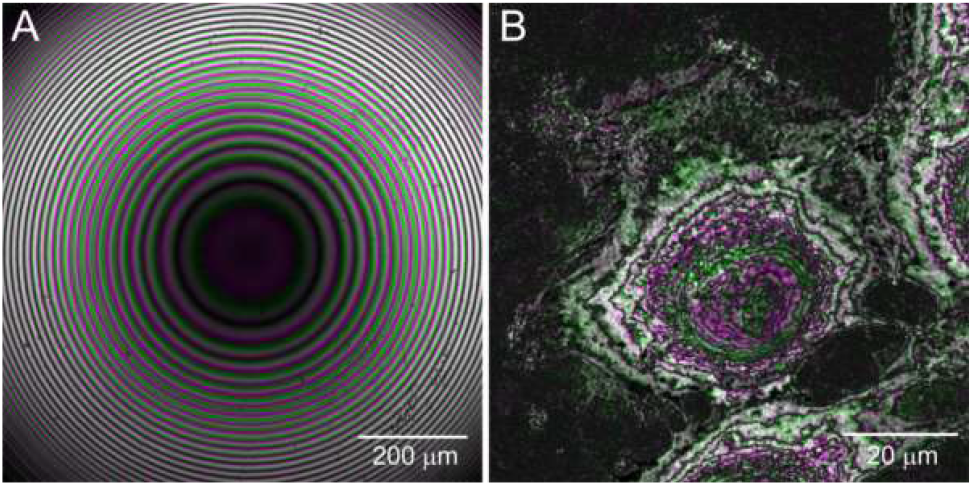
A) A false-colour composite difference image of a *f* = 72 mm lens specimen acquired using multi-wavelength IRM. The difference between |514 nm – 488 nm| is shown in green, and the difference between |543 nm – 514 nm| is shown in magenta. B) A false-colour composite difference image of fixed MeT-5A cells. The difference between |514 nm – 488 nm| is shown in green, and the difference between |543 nm – 514 nm| is shown in magenta.

Using the same radially averaged line intensity measurement method to that applied to TartanSW data, applying the difference operation to multi-wavelength IRM images of the lens resulted in FWHM antinodal fringe thicknesses of 60.0 nm for |514 nm – 488 nm| and 64.7 nm for |543 nm – 514 nm|. This offers a considerable improvement over conventional IRM where FWHM antinodal fringe thicknesses of 83.6 nm, 88.0 nm, and 108.3 nm are the thinnest possible with illumination wavelengths of 488 nm, 514 nm, and 543 nm. Further, the difference operation when applied to multi-wavelength IRM cell images showed similar improvement in antinodal fringe position precision and contrast to that obtained when applied to TartanSW datasets.

Antinodal plane thickness in SW microscopy is often attributed as equivalent to the axial resolution, but this is not strictly correct. The full theoretical structure of the widefield SW point spread function (PSF) can be obtained from the following equation [14]

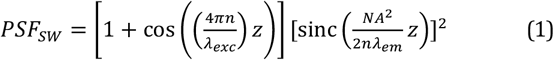

where NA is the numerical aperture of the imaging objective and _λem_ is the peak emission wavelength being detected, and the illumination intensity profile I for IRM is given by [15]

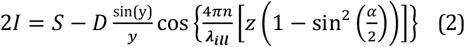

where *S* and *D* are the sum and difference of the maximum and minimum intensities respectively, *z* and *y* are axial and lateral distances, and α is half the angle of the cone of illumination. While we observe a thinning of the antinodal fringe position with the difference operation in both fluorescence and reflection interference methods studied here, this is not exactly commensurate with an improvement in axial resolution, and we have avoided this direct comparison. We also note that performing the difference operation to improve antinodal fringe thickness comes at a cost of reduced sampling density. We therefore propose the difference operation as an aid to the interpretation of multi-colour interference microscopy images, which has broad applications including surface profile analysis [16], absolute measurement of thin film thickness [17], and characterisation of protein patterns in live cells [18].

There are specific imaging conditions where the difference operation should be used with great care. For example, where there exist similar intensity values in the same region of multiple channels, such as the region close to the centre of Fig 2A, this will result in a region containing zero intensity values, as can be seen in the corresponding region of Fig. 2B. Also, since the contrast improvement is mostly observed in the higher order fringes this method is best suited to imaging thicker cell specimens with a topological shape that does not lead to overlapping antinodal planes in the axial dimension, e.g. a curved surface in contact with a flat substrate [19]. As with conventional high-resolution imaging the Nyquist sampling criterion should be fulfilled [20].

Our data show that the difference operation increases the contrast of the images, and it can increase the visibility of structures that are difficult to detect in multi-wavelength interference microscopy. This is evident in Fig. 2, where internal membrane structures that are barely visible in TartanSW data can be clearly observed after the difference operation has been applied. These internal structures can be observed even through the highly scattering cell nucleus, which we expect is represented by the dark grey region in the centre of the cell image. This increased contrast may aid in the sensitive detection of tiny deformations in the cell membrane, and in object segmentation and three-dimensional particle tracking in live cells.

## Supporting information

Visualization 1

## Funding

This work was supported by the Biotechnology and Biological Sciences Research Council, grant numbers BB/P02565X/1 and BB/T011602/1. LK was supported by the Medical Research Council and Engineering and Physical Sciences Research Council Centre for Doctoral Training in Optical Medical Imaging, grant number EP/L016559/1. PWT and GMcC were partly supported by the Medical Research Council, grant number MR/K015583/1. LMR was supported by the Leverhulme Trust.

## Disclosures

The authors declare no conflicts of interest.

See Supplement 1 for supporting content

## Notes

### Competing Interest Statement

The authors have declared no competing interest.

### Summary of Updates

Minor edits

