## Supplementary Information 1 for "A simple image processing pipeline to sharpen topology maps in multi-wavelength interference microscopy"

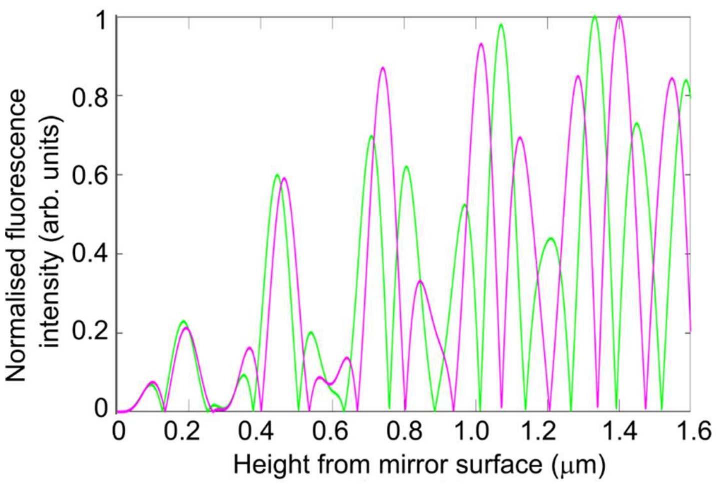


Supplementary Figure 1. Simulated axial intensity profiles resulting from difference operations between standing wave point spread functions generated using equation (1) that are propagating in air using an imaging objective lens with a numerical aperture of 0.4. The profile in magenta is the result of |550 nm -525 nm| and |525 nm – 490 nm| is shown in green.
